# Heparanase attenuates Zika virus infection by destabilizing the viral envelope protein

**DOI:** 10.1101/2025.03.16.643511

**Authors:** Jiaxin Ling, Santiago Fernández Morente, Åke Lundkvist, Jinlin Li

## Abstract

Heparanase (Hpa) is the only endoglycosidase enzyme in mammalian cells capable of cleaving heparan sulfate. In addition to its well-known functions in the regulation of glycosaminoglycans integrity, accumulating evidence indicates that Hpa plays vital roles in viral infection, while the mechanisms are not yet fully understood, especially in RNA virus infection. In this study, we report that Hpa functions as a restriction factor for Zika virus (ZIKV) infection. Our results demonstrated that Hpa, but not the enzymatic inactive mutant (Hpa-DM), resulted in degradation of the ZIKV envelope (E) protein, which could be rescued by treatment of the proteasome inhibitor (MG132) and the autophagy inhibitor (NH_4_Cl), separately. Additionally, the ubiquitination of ZIKV E did not show an significant change in the presence of Hpa. Overexpression of Hpa, but not Hpa-DM, dramatically decreased ZIKV infection in different cell models, evidenced by the reduction of viral proteins and a compromised production of infectious virions. This was further confirmed by the results in MEF cells, in which knockout Hpa enhanced ZIKV infection, while overexpression of Hpa suppressed the production of virions. In addition, ZIKV was found to downregulate the Hpa expression, which could counteract the inhibitory effects of Hpa. Altogether, our study discovers an unrecognized role of Hpa in virus infection and demonstrates that Hpa serves as a restriction factor for ZIKV infection.

**Summary:** Zika virus (ZIKV), primarily transmitted by mosquitoes, can cause a wide range of symptoms, including myalgia, fever, rash, and severe neurological complications such as microcephaly, epilepsy, and Guillain-Barré syndrome. A deep understanding of ZIKV-host cell interactions, particularly the roles of pro- and anti-viral factors, is essential for the development of effective therapeutic strategies, which are currently unavailable. In this study, we uncover that heparanase (Hpa), the only endoglycosidase enzyme in mammalian cells capable of cleaving heparan sulfate (HS), can degrade the ZIKV envelope (E) protein. Hpa exhibits an inhibitory role in ZIKV infection in different cell models. Additionally, we find that ZIKV downregulates the Hpa expression, which could be used by the virus to mitigate the inhibitory effects of Hpa. Taken together, our study demonstrates Hpa as a restriction factor in ZIKV infection and highlights complex interactions between ZIKV infection and HS machinery enzymes.

## 1. Introduction

Zika virus (ZIKV), within the genus of *Flavivirus*, was initially isolated in Uganda and has emerged as one of the most important human pathogens since the outbreak in South America in 2015 (1). ZIKV is a mosquito-borne RNA virus, which can result in various diseases ranging from non- or mild symptoms of ZIKV fever to microcephaly and other birth defects when the virus infects pregnant women (2). Adults infected by ZIKV also have a risk of developing disease, such as Guillain–Barré syndrome (3). So far, there is no approved effective drug for the prevention or treatment of ZIKV-associated diseases. Therefore, understanding the mechanisms by which ZIKV interacts with the host and thereby dissecting key factors in virus infection will favor the development of novel antiviral drugs.

Like other flaviviruses, the ZIKV’s genome consists of a single, positive-sense RNA, encoding 3 structural proteins, the capsid (C) protein, the pre-membrane (prM) protein, and the envelope (E) protein and seven non-structural proteins (NS1, NS2A, NS2B, NS3, NS4A, NS4B, and NS5) (4). After entering and releasing its genome into host cells, ZIKV initiates the translation using its positive RNA as the template to generate a single polyprotein. The synthesized polyprotein is cleaved into ten mature proteins by viral and cellular proteases (5). The nonstructural proteins mainly function as coordinators for the following ZIKV replication (6). Viral replication takes place in the replication complex (RC) formed by endoplasmic reticulum (ER) membranes, yielding a dsRNA. The intermediate negative-sense RNA will subsequently be transcribed into a positive single-stranded genomic/mRNA. The immature ZIKV particles assembled in ER will become mature infectious viruses through processing in the trans-Golgi network.

Multiple cellular factors are associated with ZIKV cell entry, including DC-SIGN, AXL, Tyro3, TIM-1, and TLR3 in different cell models (7, reviewed in 6). In addition to these functional receptors, heparan sulfate (HS) has been suggested as an important attachment factor for most flavivirus infections, although the role of HS in ZIKV infection is still controversial (7-10). HS is a member of glycosaminoglycans (GAGs), which are linear polysaccharides composed of repeating disaccharide units (11). HS is ubiquitously expressed on almost all mammalian cells’ surfaces. The wide distribution and negatively charged properties of HS make it a co-factor for the cell entry of various viruses (10).

Physiologically, heparanase (Hpa) is the only endoglycosidase enzyme responsible for cleaving HS on the surface of mammalian cells (12, 13). Hpa is well known for its roles in tumor metastasis (14). Due to the regulatory roles in cleaving HS and the tight relationship between HS and viruses, HS-binding viruses have been reported to hijack the heparanase for promoting viral release from cells (15-17). Herpes simplex virus (HSV-1) upregulates the expression of Hpa to facilitate the release of the mature virion by enchaining the shedding of syndecan-1 or reducing the HS on the cell surface (16, 17). Porcine Reproductive and Respiratory Syndrome Virus (PRRSV) also uses a similar strategy by increasing the expression of Hpa to favor its release (15). Beyond its enzymatic roles in regulating HS, Hpa was recently found to remodel the host environment for pathogens invasion by dysregulating the innate defense responses including the interferon and DNA damage response signal pathways (18, 19), indicating that Hpa might have multiple crucial functions in virus infection and pathogenesis. However, there is limited knowledge available regarding the roles of Hpa in flavivirus infections. In this study, we demonstrate that Hpa functions as a host restriction factor to inhibit ZIKV infection by affecting the stability of viral protein.

## 2. Materials and Methods

### 2.1 Antibodies and reagents

The following antibodies were used in this study: Monoclonal ANTI-FLAG® M2 antibody (1:5000, Sigma, F1804); Polyclonal anti-heparanase S63 (1:1000, 1:200IF) was kindly provided by Prof. Vlodavsky (Technion Integrated Cancer Center, Haifa, Israel); Ubiquitin Antibody (P4D1) (1:1000, Santa Cruz, sc-8017); GAPDH Monoclonal antibody (1:10000, Proteintech, 60004-1-Ig); Zika virus Envelope protein antibody (1:2000, Gene Tex, GTX133314); Goat anti-Mouse IgG (H+L) Secondary Antibody, HRP (Invitrogen, 31430); Goat anti-Rabbit IgG (H+L) Cross-Adsorbed Secondary Antibody, HRP (Invitrogen, G21234), Goat anti-Rabbit IgG (H+L) Cross-Adsorbed Secondary Antibody, Alexa Fluor™ 594 (Invitrogen, A-11012). Reagents in this study: G418 solution (Merck, 4727878001); Puromycin Dihydrochloride (Gibco, A1113803); Lipofectamine 3000 (Invitrogen, L3000015). MG132 (MCE, HY-13259).

### 2.2. Virus and Plasmids

Zika virus (strain MR-766) was propagated in Vero E6 cells. The titers of ZIKV were determined by plaque assay as described below. The pcDNA3 plasmids expressing wild-type heparanase (pcDNA3-Hpa) or inactive enzymatic mutant (pcDNA3-Hpa-DM) as well as empty vector are the gifts from Prof. Israel Vlodavsky (Technion Integrated Cancer Center, Haifa, Israel) (20). The plasmids expressing ZIKV envelope protein or NS1 protein were constructed by amplifying the coding sequence and cloned into between PstI and XhoI restriction sites in the pCMV-3Tag plasmid. The CDS of Hpa or Hpa (DM) was amplified by the primers (5’-<u>CCGCTCGAG</u>ATGCTGCTGCGCTCGAAG-3’ and 5’-<u>TGCTCTAGA</u>TCAGATGCAAGCAGCAACTTT-3’) and cloned in the Xhol and Xbal sites of PLVX-IRES-Puro Vector (Clontech, TaKaRa) to construct FLVX-Hpa or FLVX-Hpa (DM) for generating an Hpa or Hpa (DM) stably expressing cell line.

### 2.3 Cell culture

HEK293T (ATCC: CRL-3216), Vero E6 cells (ATCC: CRL-1586) and Huh7 (ATCC: PTA-4583) were from American Type Culture Collection. Mouse embryonic fibroblasts (MEF) of wild type (MEF), Hpa knockout (MEF-Hpa-KO), and Hpa overexpression (MEF-Hpa) (21, 22) were kindly provided by Prof. Jin-ping Li at Uppsala University. Cells were cultured in Dulbecco’s modified Eagle medium (DMEM, Gibco, 10569010) supplemented with 10% fetal bovine serum (FBS) (Gibco, A5256701), and penicillin-streptomycin (Gibco, 15140122) at 50 units/ml. Cells were incubated at 37 °C in a 5% CO_2_ incubator. For Hpa / Hpa-DM-overexpressing Huh7 or HEK-293 cells, G418 (600 μg/mL) or Puromycin (2 µg/mL) were added to the medium to select positive cells containing Hpa or Hpa mutant (Hpa-DM) expression separately.

### 2.4. Construction of cell lines stably expressing heparanase and its inactive enzymatic mutant

The pcDNA3-Hpa, pcDNA3-Hpa-DM, or pcDNA3-vec plasmids were transfected to Huh7 cells separately by lipofectamine 3000 transfection reagent (Invitrogen, L3000015) and selected under G418 (600 µg/ml, Merck, 4727878001). The dead cells were removed by replacing with the fresh media containing 600 µg/mL G418 and the culture was kept for at least two weeks. To construct the HEK-293 cell stably expressing Hpa or Hpa-DM, lentivirus was packaged by co-transfection of FLVX-EXT1 with the 2nd generation lentiviral systems plasmids psPAX2 and pMD2.G (a gift from Didier Trono, Addgene plasmid #12260 and # 12259). Lentivirus-transduced HEK-293 cells were selected under 2 µg/mL puromycin for at least 2 weeks. The expression of Hpa was examined by western blot using an antibody (S63) against Hpa.

### 2.5. Plaque assay

Supernatants containing infectious ZIKV was collected and stored at −80°C before measuring the titers. For plaque assay analyses, Vero E6 cells were seeded into 24-well plates and grown to monolayer. Serial dilutions of viral stocks were incubated with cells for 1 h at 37 °C, followed by adding 1 ml 0.8% agar media (Sigma-Aldrich, A5431) to each well. After 48 to 72 h, cells were fixed and stained with 0.5% Crystal Violet Solution (Sigma-Aldrich, V5265), and the number of plaques was counted.

### 2.6. Western blots and immunoprecipitation

Cells were lysed in RIPA buffer (25 mM Tris•HCl pH 7.6, 150 mM NaCl, 1% NP-40, 1% sodium deoxycholate, 0.1% SDS). Protease Inhibitor Cocktail (Thermo Fisher Scientific, 78430) was supplemented into lysis buffer to prevent proteolytic degradation of cell lysis. Laemmli SDS sample buffer (Bio-Rad, 1610747) was added to each sample, followed by boiling for 10 min at 100°C. The cell lysates were fractionated by 10% or 12% PAGE gel, transferred to the PVDF membrane using tank transfer, and probed with the primary antibody, and then followed by the secondary antibody. The immunocomplexes were visualized by SuperSignal™ West Pico PLUS Chemiluminescent Substrate (Thermo Fisher Scientific, 34580) using ChemiDoc MP Imaging System (Bio-Rad). For detecting the ubiquitination of ZIKV E protein, the cells were lysed on ice for 30 min under the denaturing condition with lysis buffer (50 mM Tris-HCl pH 7.4, 150 mM NaCl, 1% Triton X-100, 1mM EDTA, 10% glycerol, 20 mM NEM, 2 mM Iodoacetamide, protease inhibitors cocktail) supplemented with 1% SDS, followed by dilution to 0.1% SDS before the immunoprecipitation using ANTI-FLAG® M2 Affinity Gel (A2220; Sigma) at 4°C for three hours. After 3 times washing with lysis buffer and one time washing by TBS (pH=7.4) supplemented with protease inhibitor, FLAG peptide (F4799; Sigma) was added to elute the immunocomplexes by incubating at 4°C for 30 min.

### 2.7. Real-time PCR

RNA was isolated from cell pellets by RNeasy Mini Kits (Qiagen, 74104) according to the protocol from the kit. One microgram RNA was used to prepare cDNA by using the iScript™ cDNA Synthesis Kit (Bio-Rad, 1708891) based on the protocol: 25°C for 5 mins, 46°C for 20 mins, and 95°C for 1 min. One microliter of reaction was utilized to run the following real-time PCR using SYBR® Green Master Mix (Bio-Rad, 1725271) with cycling conditions: 95°C for 30 sec, followed by 40 cycles of denaturation at 95°C for 10 sec and annealing/extension at 60°C for 60 sec. Fold change was calculated as: Fold Change = 2^-Δ(ΔCt)^. The primers used in this study for quantitative PCR are listed below. ZIKV E gene (F-5’TTGGTCATGATACTGCTGATTGC3’; R-5’CCTTCCACAAAGTCCCTATTGC3’); human GAPDH gene (F-5’TGGGCTACACTGAGCACCAG3’; R-5’ GGGTGTCGCTGTTGAAGTC3’); human GRP78 (F-5’CATCACGCCGTCCTATGTCG’3; R-5’CGTCAAAGACCGTGTTCTCG’3) and human HPSE (F-5’ACGGCTAAGATGCTGAAGAGC3’; R-5’TTCCTTGGTAGCAGTCCGTG3’).

## 3. Results

### 3.1. Heparanase downregulates the expression of the ZIKV envelope

ZIKV E protein plays multiple roles in ZIKV infection and pathogenesis. Any alteration in ZIKV E protein expression may dramatically remodel viral infection. To investigate whether Hpa has any effect on ZIKV E protein, FLAG-tagged E (FLAG-E) and Hpa or Hpa-DM (the enzymatic inactive mutant of Hpa) plasmids were co-transfected into HEK-293 cells and the expression of proteins was assessed. Hpa, but not Hpa-DM, significantly downregulated the expression of FLAG-E in co-transfected cells, suggesting that Hpa can mediate the downregulation of FLAG-E expression independently of ZIKV replication (Fig. 1A, 1B). To further provide evidence for the specific inhibitory role of Hpa on ZIKV E protein and rule out the possibility of promoter competition, we cloned the CDS of a non-structural protein of ZIKV, NS1, into the same plasmid vector as the one used to express ZIKV E protein. In the same experimental set, the expression of FLAG-NS1 did not change when overexpressing Hpa or Hpa-DM (Fig. 1C). Collectively, the results suggest Hpa can result in the downregulation of ZIKV E protein.

**Figure 1.**
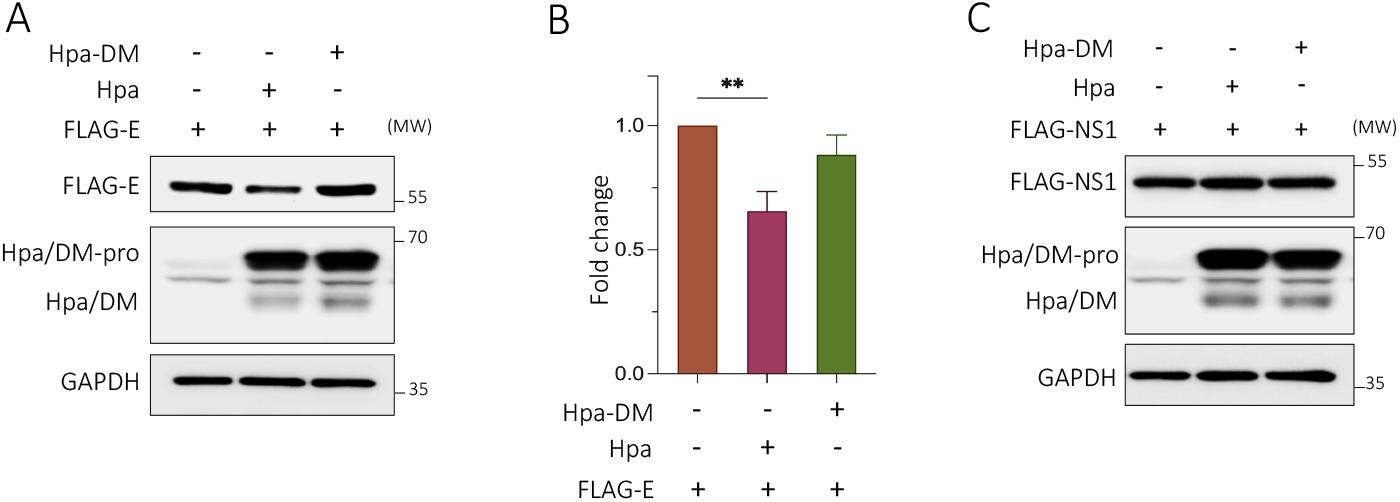
Heparanase downregulates the expression of the ZIKV envelope. (**A**) FLAG-E was co-transfected with Hpa, Hpa-DM, or vector to HEK-293 cells separately. The expression of ZIKV E protein and Hpa or Hpa-DM were examined by western blot. One representative experiment of three was shown. (**B**) The intensity of ZIKV E protein and GAPDH specific bands were quantified using Image J software. The relative intensity of the envelope was calculated as the ratio of FLAG-E and GAPDH densitometry values. The data were presented as fold change by normalizing to the relative intensity of FLAG-E in the vector control condition. Mean ± SD from three experiments was calculated. Statistical analysis was performed using Student’s t-test. **p≤ 0.01. (**C**) FLAG-NS1 was co-transfected with Hpa, Hpa-DM, or vector to HEK-293 cells separately. The expression of NS1 protein and Hpa or Hpa-DM was assessed by western blot.

### 3.2. Hpa-mediated ZIKV E protein degradation is via the lysosomal proteolysis

Next, we endeavored to understand how Hpa resulted in the downregulation of ZIKV E protein. The ubiquitination-proteasome system and autophagy are the two main protein degradation machinery in mammalian cells that regulate protein homeostasis.

We utilized MG132, a proteasome inhibitor or ammonium chloride (NH_4_Cl), a lysosomal inhibitor that raises the lysosomal pH to inhibit the autophagy-mediated protein degradation, to test the possible pathways involved in Hpa-mediated ZIKV E protein degradation. After co-transfection of FLAG-E and Hpa, MG132 was added to the cells at the last 8 h before harvest. As demonstrated above, Hpa significantly downregulated the expression of FLAG-E. However, the decrease was markedly attenuated when treated with MG132 (Fig. 2A, 2B). Similarly, the treatment of NH_4_Cl also resulted in the restoration of the FLAG-E level in the presence of Hpa (Fig. 2C, 2D). Proteasome-mediated degradation usually involves the ubiquitination of the target protein. We found that FLAG-E could be ubiquitinated and the ubiquitination of FLAG-E did not exhibit a significant increase in the presence of Hpa (Fig. 2E). Altogether, our results suggest that Hpa-mediated ZIKV E protein downregulation is via the lysosomal proteolysis instead of the proteasome-mediated degradation pathway.

**Figure 2.**
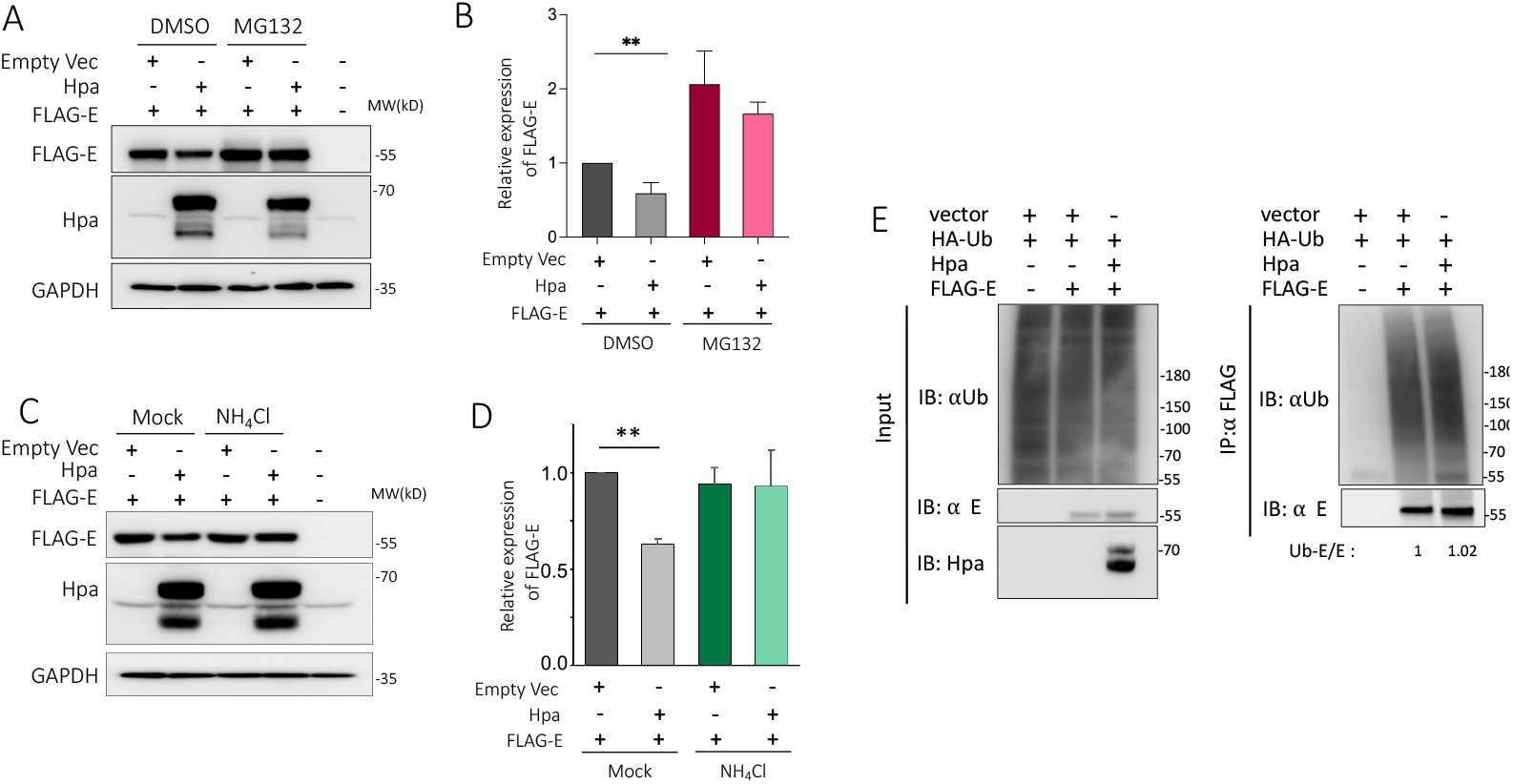
Hpa-mediated ZIKV E protein degradation is via lysosomal proteolysis. FLAG-E was co-transfected with Hpa or vector. After 40 h, the cells were treated with or without MG132 (**A**) or NH_4_Cl (**C**) for 8h. The expression of FLAG-E and Hpa were evaluated by western blot using anti-FLAG or anti-Hpa antibodies separately. One experiment of three was shown here. (**B**) or (**D**) The intensity of ZIKV E protein and GAPDH specific bands in (A) or (C) were quantified using Image J. The relative expression of FLAG-E was calculated by the intensity ratio of FLAG-E / GAPDH and normalized to the value in the co-transfection of FLAG-E with vector without treatment. The data was shown as mean ± SD from three experiments. Statistical analysis was performed using Student’s t-test. **p≤ 0.01. (**E**) HA-Ub was co-transfected with FLAG-E and untagged Hpa. Cells were treated with MG132 (10µM) for 6h before the immunoprecipitation with anti-FLAG beads in the denature conditions and the blots were probed with an anti-Ub antibody. The intensity of ZIKV E Ub conjugates was normalized by the intensity of the ZIKV E specific bands separately.

### 3.3. Heparanase inhibits ZIKV infection

After demonstrating that Hpa enhanced the degradation of ZIKV E protein, we further examined whether Hpa remodeled ZIKV infection by using different Hpa knockout and overexpression cell models. We constructed Huh 7 stably overexpressing Hpa (Huh7-Hpa) or a Hpa enzymatic inactive mutant (Huh7-Hpa-DM) cell lines. Both Hpa and Hpa-DM were expressed at a comparable level in the Huh7-Hpa and Huh7-Hpa-DM cells (Fig. S1A, S1B). Huh7-Hpa or Huh7-Hpa-DM were infected by ZIKV and the expression of ZIKV E protein was assessed to monitor the ZIKV infection. Compared to the Huh7-Vec control cells, ZIKV E protein expression was reduced in Huh7-Hpa cells. However, there is only a marginal effect on the expression of ZIKV E in Huh7-DM cells (Fig. 3A). This conclusion was further supported by the similar results from HEK-293 Hpa wildtype and overexpression cells infected by ZIKV (Fig. 3B). To further corroborate the inhibitory role of Hpa in ZIKV infection, we infected MEF cells, Hpa overexpressing MEF cells (MEF-Hpa) or Hpa-knockout MEF cells (MEF-Hpa-KO) with ZIKV. In line with the data from Huh7 and HEK293, a significant decrease was observed in MEF-Hpa cells, while knocking out Hpa enhanced the expression of E in MEF-Hpa-KO cells (Fig. 3C). We also checked the replication of ZIKV by qPCR using the specific primers targeting the ZIKV E gene, overexpression of Hpa did not downregulate ZIKV replication (Fig. 3D). As expected, the infectious viruses produced by the Huh7-Hpa and MEF-Hpa cell lines were markedly lower than those produced by the Huh7-Vec and MEF cells, respectively (Fig. 3E, 3F). While this is not the case for the inactivated Hpa overexpressing cell line, Huh7-DM, which released a quite similar amount of virus as in the vector control cells (Fig. 3E). Knocking out Hpa enhanced the ZIKV particle production in MEF cells (Fig. 3F). The data from various cell models indicates that Hpa impedes ZIKV infection independent of cell types.

**Figure 3.**
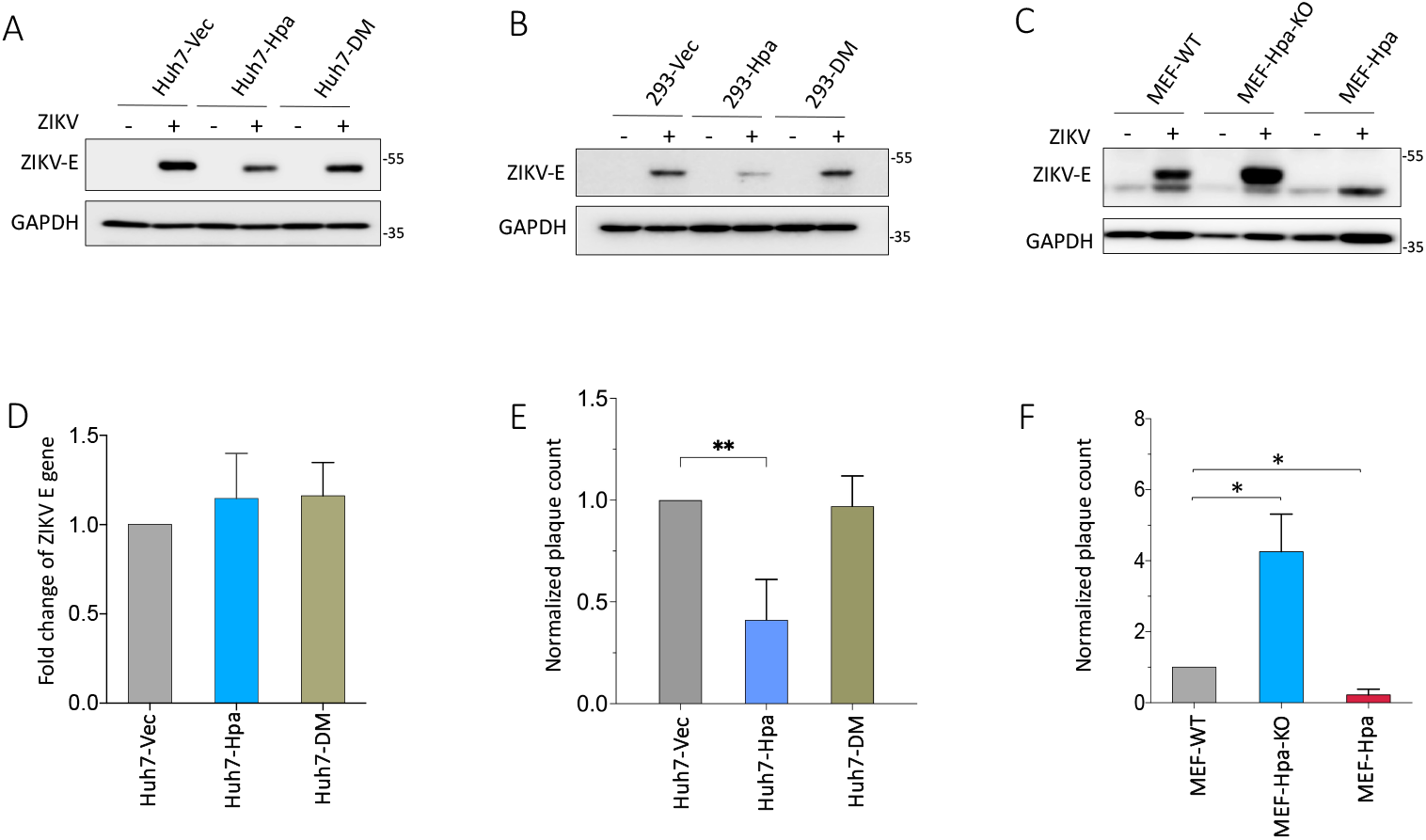
Heparanase inhibits ZIKV infection. (**A**) Huh7 or (**B**) HEK-293 stably overexpressing Hpa or Hpa-DM cell lines were infected by ZIKV (MOI=0.1). After 48h (for Huh7) and 24 h (for HEK-293), the expression of E protein was assessed by western blot using the specific antibody. (**C**) MEF or MEF-Hpa-KO cells were infected with ZIKV (MOI=0.1). The expression of ZIKV E protein was assessed by western blot at 48h post-infection. (**D**) The replication of ZIKV was evaluated by qPCR using the specific primers targeting the ZIKV E gene. The results were presented as fold change relative to the infection of Huh7-Vec cells and the data were shown as mean ± SD from two experiments. (**E**) or (**F**) The titer of virus particles released into the supernatant in (**A**) or (**C**) was measured by standard plaque assay. The number of plaques was normalized by vector control or wild-type cells and was expressed as mean ± SD of two or three experiments. Statistical analysis was performed using Student’s t-test. *p≤ 0.05, **p≤ 0.01.

### 3.4. ZIKV infection attenuates heparanase expression

In order to create a friendly environment for viral replication and propagation, viruses have evolved with different strategies to cope with those restriction factors from the host cells. In the last set of experiments, we aimed to dissect the effects of ZIKV infection on Hpa by monitoring the expression of Hpa at different time points post-ZIKV infection in Huh7 cells. The expression of the ZIKV E protein was utilized to indicate the ZIKV infection. The ZIKV E protein started to be detected at 24 h and was dramatically increased at 48h post-infection (Fig. 4A). The samples from each time point were applied to examine the Hpa expression. Hpa mRNA did not alter at the early time point (4h) after ZIKV infection, while it slightly increased at 12 h post-infection and was followed by a gradual downregulation from 12h to 48h after infection (Fig. 4B). A similar pattern of Hpa mRNA expression changes was also observed in HEK293 cells infected with ZIKV. (Figure S2). The significant decrease of Hpa at the 48h post-infection is related to the late phase of ZIKV infection with the extensive expression of structural proteins (Fig. 4). To rule out the possibility that the downregulation of Hpa mRNA at 48h after ZIKV infection was non-specific, we examined the mRNA expression of another cellular gene, GRP78, at 48h post-ZIKV infection in Huh7. As shown in Fig. 4C, the GRP78 mRNA expression was upregulated. It is likely that due to the low expression of endogenous Hpa, we failed to detect the Hpa in Huh 7 cells by western blot, even when using different available antibodies against Hpa. The data indicated that Hpa was downregulated in the late phase of ZIKV infection.

**Figure 4.**
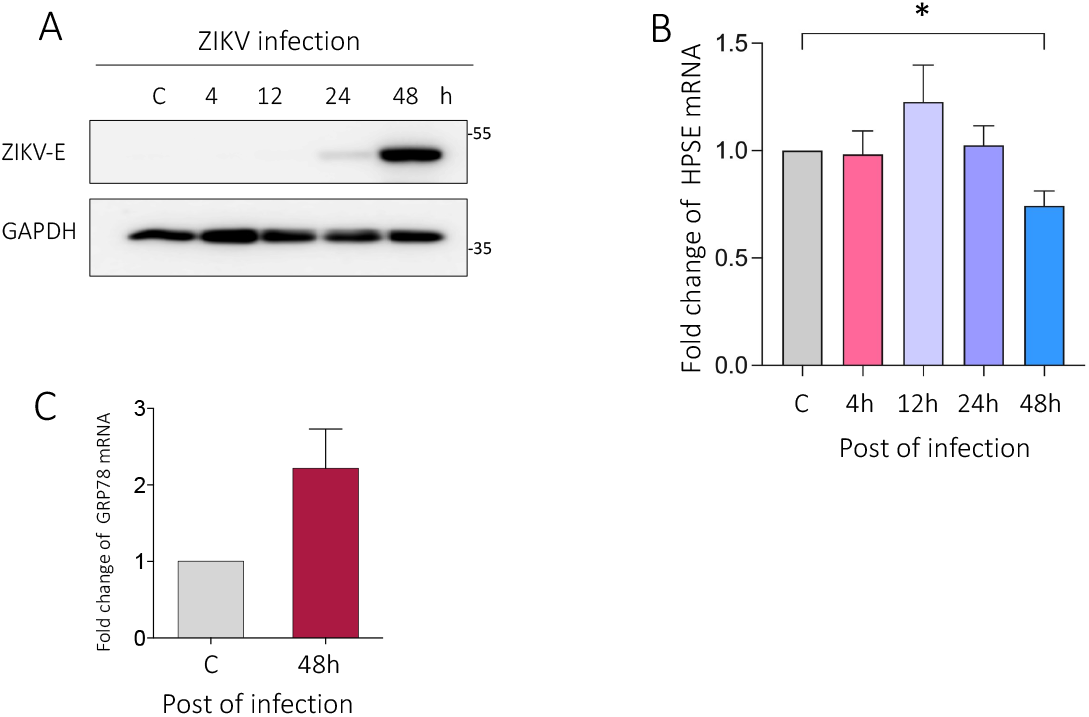
ZIKV infection attenuates heparanase expression. (**A**) Huh7 cells were infected by ZIKV (MOI=0.1). Cells were harvested at different time points after infection as shown in the figure. The expression of ZIKV E protein was examined by western blot to monitor ZIKV infection. (**B**) The expression of HPSE at the mRNA level was quantified by qPCR. The change of HPSE mRNA was calculated as fold change related to the mock infection and results were shown as mean ± SD from three experiments. Statistical analyses were performed by one-way ANOVA. *p≤ 0.05. (**C**) The expression of GRP78 at the mRNA level was evaluated by qPCR at 48h post-ZIKV infection from the samples in (B).

## 4. Discussion

Although the functions of Hpa on tumor metastasis, angiogenesis, and inflammation have been well documented, very little is known about its roles in RNA virus infections, especially for its contributions to virus infection beyond its impacts on HS cleavage. In this study, we uncovered an unrecognized role of Hpa in RNA virus infection and demonstrated that Hpa promoted the degradation of ZIKV E protein, by which to inhibit ZIKV infection. To overcome the restriction effects from Hpa, ZIKV downregulated the expression of Hpa to fulfill its production successfully. Our data points out that Hpa is a restriction factor for ZIKV infection.

HS is commonly utilized by flaviviruses as an attachment factor. Different from other members of flavivirus, such as DENV, TBEV, and JEV where the role of HS as co-receptors has been established, the function of HS in ZIKV entry is still conflicting from different studies (8, 9, 23, 24). However, accumulating evidence from our and other studies suggested that ZIKV did not exploit HS to enter cells (25, 26). Despite the high binding affinity to HS is beneficial for viral cell attachment, the viruses face the challenge of evading the HS trap when they release. To cope with this, some evolutionary tactics are hijacked by viruses, such as the upregulation of Hpa by HSV-1 and PRRSV during their life cycles (15, 17). The independent relationship between HS and ZIKV may shape viral transmission pathways from cell to cell. Among the flaviviruses, ZIKV is the only member that is closely associated with congenital infection in humans (27). So far, DENV and WNV, two other flaviviruses whose cell entry is HS-dependent, have been reported to upregulate Hpa during the infections (10, 28). In our study, we showed the expression of Hpa was downregulated in the late stage of ZIKV infection, which, in some sense, agrees with the scenario that ZIKV does not need to overcome the HS trap when it is released by upregulating Hpa. Whether the interactions between ZIKV and cellular HS machinery (including HS and other enzymes involved in the biosynthesis and regulation of HS, such as Hpa, Exostosin Glycosyltransferase (EXT), and N-deacetylase and N-sulfotransferase (NDST)) impacts ZIKV pathogenesis requires further investigation.

Another important discovery in the current study is that we found that Hpa degraded the ZIKV E protein. The ubiquitin-proteasome pathway and lysosomal proteolysis are the two major systems for protein degradation (29). As a proteasome inhibitor, MG132 also inhibits some lysosomal hydrolases (30). In our study, the restoration of Hpa-mediated downregulation of ZIKV E by MG132 or NH_4_Cl and the unchanged ubiquitination of ZIKV E protein in the presence of Hpa, point out that the degradation of E is probably through lysosomal proteolysis. Autophagy is a cellular degradation pathway mediated by lysosomes. There are three different kinds of autophagy based on its cargo delivery manner and membrane dynamics: macroautophagy, microautophagy, and chaperon-mediated autophagy (CMA) (31). In contrast to macroautophagy and microautophagy that require membrane deformation, CMA-mediated degradation initiates with the recognition of the KFERQ-like motif on the targeted protein by molecular chaperone HSC70, followed by its transport into the lysosome with the assistance of the lysosomal membrane protein LAMP2A (32). Though no classic KFERQ-like motif was found on the ZIKV E protein, the advanced motifs, **NRDFV** (asparagine motif) and **KRTLV** (acetylation and phosphorylation-activated motifs) were mapped (33), which might be also recognized by HSC70 and be subjected to the CMA-mediated degradation. Since all three categories of autophagy can selectively degrade proteins, which one (s) is involved in the Hpa-mediated ZIKV E protein degradation needs to be clarified in future studies.

Hpa has been reported to promote autophagy *in vivo* and *in vitro*, and the enhanced autophagy has been ascribed to the downregulation of phospho-RPS6KB/p70 S6-kinase, a substrate of rapamycin complex 1(MTORC1) (34). Though the results are controversial regarding the role of autophagy in ZIKV infection, many studies have shown that ZIKV infection induced autophagy. In the murine model, ZIKV infection caused the upregulation of autophagy in mouse neural tissue (35). The formation of autophagosomes was induced in the skin fibroblasts following ZIKV infection (36). ZIKV infection also activates autophagy in human trophoblast cells and mouse placenta (37). In our study, we observed a significant decrease in Hpa expression at the late time point of ZIKV infection. Taking account into the positive regulation of Hpa on autophagy, our data indicate that ZIKV infection might downregulate autophagy at a late time point of ZIKV infection, which agrees with a recent study that found autophagy was suppressed in the late phase of ZIKV infection (38). Altogether, our study points out a restricted role of Hpa in ZIKV infection and highlights complex interactions among ZIKV infection, Hpa, and autophagy.

## Supplementary Materials

The following supporting information can be downloaded at: Figure S1: The expression of Hpa and Hpa-DM in the Huh7 cells and Figure S2: The expression of HPSE mRNA in the HEK293 cells infected by ZIKV.

## Author Contributions

Conceptualization, J.Li and J.Ling; methodology, S.F.M., J.Li, J.Ling; validation, S.F.M.,and J.Ling; formal analysis, J.Li and J.Ling; investigation, S.F.M.,J.Ling and J.Li; resources, Å.L. and J.Li; data curation, S.F.M., J.Li; writing— original draft preparation, J.Li and J.Ling; writing—review and editing, J.Ling, S.F.M., Å.L. and J.Li; visualization, J.Li; supervision, J.Li and J.Ling; funding acquisition, Å.L, J.Ling and J.Li. All authors have read and agreed to the published version of the manuscript.

## Data Availability Statement

The data that support the findings of this study are available from the corresponding author upon reasonable request.

## Acknowledgments

We sincerely thank Prof. Israel Vlodavsky from Technion Integrated Cancer Center, Haifa, Israel and Prof. Jin-ping Li from Uppsala University, Sweden, for sharing the antibodies, plasmids and cell lines.

