## Supplementary material for "Heparanase attenuates Zika virus infection by destabilizing the viral envelope protein": Fig.S1 and Fig S2

### Supplementary Materials

#### Methods and materials

##### Immunofluorescence assay

Huh7 cells stably overexpressing wide-type heparanase (Hpa) or the mutant (Hpa-DM) were grown on coverslips for around 24 h. For immunofluorescence analysis, the cells were fixed with cold methanol for 20 min at -20°C, followed by 3 times washing with PBS and each time 5 min. The fixed cells were permeabilized with 0.05% TritonX-100 in PBS and blocked with 4% bovine serum albumin for 40 min. After incubation with primary antibodies at room temperature (RT) for 1 h, the cells were washed with PBS 4 times with 5 min each time. Then, appropriate Alexa Fluor-conjugated secondary antibodies were applied to incubate with cells at RT for 1 h in a dark incubation box, followed by 4 times washing and mounting in Vectashield-containing DAPI (Vector Laboratories, Inc. Burlingame, CA, USA). The images were acquired by fluorescence microscope (Eclipse 90i, Nikon, Tokyo, Japan). Analysis of images was performed by using the ImageJ software.

Fig.S1

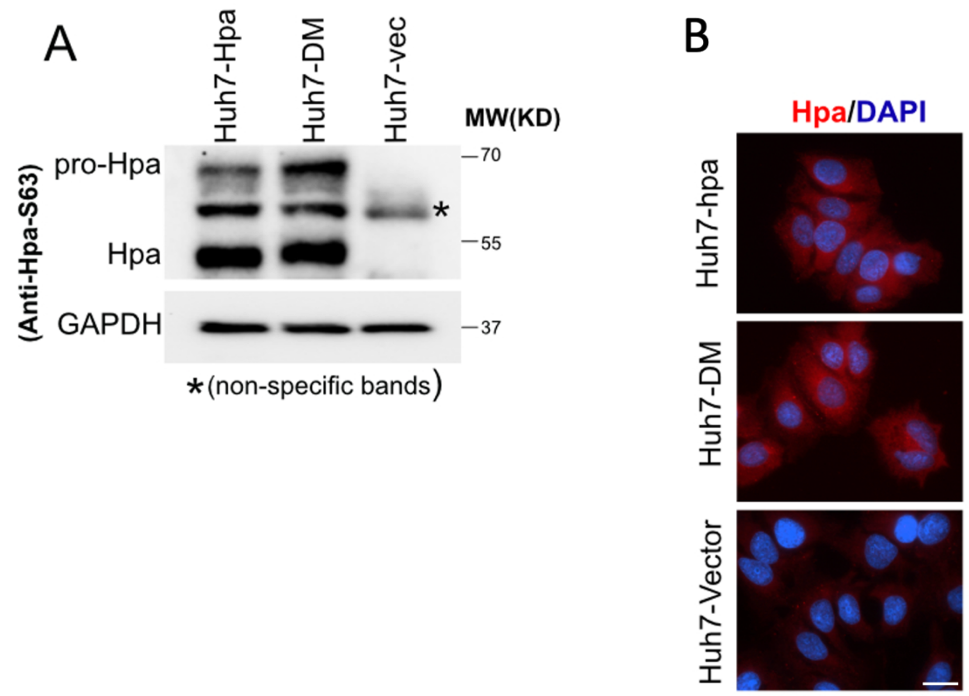


**Figure S1. The expression of Hpa and Hpa-DM in the Huh7 cells**. The expression of Hpa or Hpa-DM in the huh7 stably overexpressing Hpa or Hpa-DM cell lines was assessed by western blot (**A**) and by immunofluorescence assay (**B**) separately. Scale bar = 10 μm.

Fig.S2

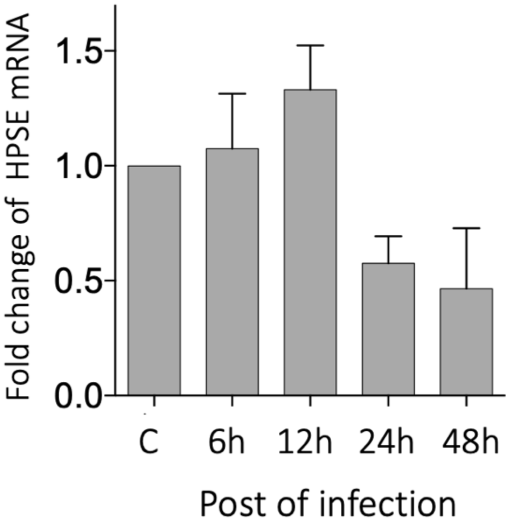


**Figure S2. The expression of HPSE mRNA in the HEK293 cells infected by ZIKV.** The expression of HPSE at the mRNA level was quantified by real-time PCR. The change of HPSE mRNA was calculated as fold change related to the mock infection and results were shown as mean ± SD from two experiments.
